# Visualizing cassava bacterial blight at the molecular level using CRISPR-mediated homology-directed repair

**DOI:** 10.1101/2020.05.14.090928

**Authors:** Kira M. Veley, Ihuoma Okwuonu, Greg Jensen, Marisa Yoder, Nigel J. Taylor, Blake C. Meyers, Rebecca S. Bart

## Abstract

Research on a few model, plant-pathogen systems has benefitted from years of tool and resource development. This is not the case for the vast majority of economically and nutritionally important plants, creating a crop improvement bottleneck. Cassava bacterial blight (CBB), caused by *Xanthomonas axonopodis* pv. *manihotis* (*Xam*), is an important disease in all regions where cassava (*Manihot esculenta* Crantz) is grown. Here we describe development of a tool for molecular-level visualization of CBB dynamics *in vivo*. Using CRISPR-mediated homology-directed repair (HDR), we generated plants containing scarless insertion of a GFP reporter at the CBB susceptibility (*S*) gene *MeSWEET10a*. Activation of *MeSWEET10a-GFP* by *Xam* was subsequently visualized at transcriptional and translational levels. Development of this tool was facilitated by a time-saving, adaptable strategy for identifying successful products of HDR, currently a limiting factor in plant research. This strategy has the potential to enable such research in other systems, improving the practicality of HDR-based experimentation.

## INTRODUCTION

Many plant pathogens, including bacteria, fungi, oomycetes, and animals, secrete proteins (i.e. effectors) into host cells to modulate host defense responses and alter gene expression to benefit the pathogen (Hogenhout *et al*., 2009). Understanding how effectors facilitate pathogenicity at the molecular level is critical for improving resistance and disrupting the processes underlying host colonization and disease. For many bacterial pathogens, effectors play an important role in disease, by targeting and altering specific host molecules (e.g. proteins, RNA, DNA) to benefit the bacteria (Feng and Zhou, 2012). Some such interactions are recognized by plant resistance (*R*) genes, resulting in an immune response. A current strategy for improving crop disease tolerance is the introduction of *R* genes into susceptible varieties (Dangl *et al*., 2013). Though effective, this strategy requires the existence or engineering of active *R* genes. Even if a suitable *R* gene is available, this strategy may be overcome by the pathogen (Brown, 2015).

Targeting susceptibility (*S*) genes is a viable, and potentially more robust alternative for engineering disease resistance in plants (Xu *et al*., 2019). *S* genes are typically necessary for normal plant growth and development, but present liabilities within the host’s genome which can be exploited by the pathogen to promote virulence (van Schie and Takken, 2014). Preventing such exploitation is a goal for engineering resistance. As with *R* genes, this strategy also requires in-depth, molecular-level knowledge of the relationship between the pathogen and host cells. Most well-characterized *S* gene – effector relationships are described from the model bacterial pathogens *Pseudomonas* spp. and *Xanthomonas* spp. (Büttner, 2016; Khan *et al*., 2018), which secrete a variety of effector proteins into host plant cells via the type III secretion system (T3SS). Canonical transcription activator-like (TAL) effectors are common in *Xanthomonas* spp. and serve as model effectors for better understanding disease susceptibility and transcriptional activation (Antony *et al*., 2010; Cernadas *et al*., 2014; Cohn *et al*., 2014; Cox *et al*., 2017; Hu *et al*., 2014; Phillips *et al*., 2017; Verdier *et al*., 2012). Each TAL effector activates transcription by recognizing and binding a specific, largely predictable sequence (effector binding element, EBE) present in the promoter of a targeted *S* gene (Boch *et al*., 2009). Genes encoding the SWEET (<u>S</u>ugars <u>W</u>ill <u>E</u>ventually be <u>E</u>xported <u>T</u>ransporters) sugar transporters from rice (*Oryza sativa* L.) are some of the best characterized *S* genes targeted by *Xanthomonas* (Chen *et al*., 2010; Yuan and Wang, 2013). The rice genome contains 22 *SWEET* genes, at least five of which (*OsSWEET11-15*) are *S* genes, playing roles in promoting susceptibility to rice bacterial blight, caused by *X. oryzae* pv. *oryzae* (*Xoo*) (Streubel *et al*., 2013). *Xoo* TAL effectors PthXo1 and PthXo2 activate transcription of *OsSWEET11* and *OsSWEET13*, respectively, while *OsSWEET14* is targeted by at least four TALs (AvrXa7, PthXo3, TalC, and TalF) (Doucouré *et al*., 2018; Streubel *et al*., 2013; Tran *et al*., 2018; Yang *et al*., 2006; Yang and White, 2004; Yu *et al*., 2011; Yuan *et al*., 2011; Zhou *et al*., 2015). The effectors targeting the *S* genes *OsSWEET12* and *OsSWEET15* have yet to be determined (Xu *et al*., 2019). Naturally occurring mutations in the EBEs of *OsSWEET11*, *OsSWEET13*, and *OsSWEET14* confer resistance to *Xoo* by preventing their transcriptional activation (Chu *et al*., 2006; Hutin *et al*., 2015; Liu *et al*., 2011; Yang *et al*., 2006; Yuan *et al*., 2009; Zaka *et al*., 2018; Zhou *et al*., 2015). Recently, the use of TALEN and CRISPR-Cas gene editing techniques to edit *SWEET* and other *S* genes has been reported to improve disease resistance in rice (Blanvillain-Baufumé *et al*., 2017; Li *et al*., 2012; Oliva *et al*., 2019; Wang *et al*., 2016; Xu *et al*., 2019).

Modifications to SWEET genes and other types of *S* genes have also been successful for improving resistance in pathosystems of citrus (Peng *et al*., 2017), cucumber (Chandrasekaran *et al*., 2016), tomato (Nekrasov *et al*., 2017), wheat (Wang *et al*., 2014; Zhang *et al*., 2017) and cassava (Gomez *et al*., 2019).

*Xanthomonas axonopodis* pv. *manihotis* (*Xam*), causes cassava bacterial blight (CBB). CBB is a worldwide problem, occurring in all regions where cassava is grown. Severity is extremely variable and unpredictable, sometimes resulting in 100% crop loss (Lin *et al*., 2019; López and Bernal, 2012). No genetic resistance to CBB is available. Symptoms of an infection begin with dark green lesions on leaves (i.e. water soaking), followed by leaf wilting and eventual death of aboveground tissues. Upon infection, *Xam* delivers 20 to 30 effectors via the T3SS into cassava cells, including five TAL effectors (Bart *et al*., 2012). While the majority of host gene targets have yet to be identified, TAL14 and TAL20 are known to promote virulence by inducing *S* genes encoding two pectate lyases and a SWEET family sugar transporter, MeSWEET10a, respectively (Cohn *et al*., 2014, 2016). Most importantly, ectopic induction of *MeSWEET10a* in infected cassava leaves is required for the water soaking phenotype and full pathogen virulence (Cohn *et al*., 2014).

Understanding how bacterial pathogens manipulate host genes to promote disease at the molecular level is essential for engineering robust, disease resistant plants. Despite the observed role for *MeSWEET10a* in susceptibility, it is currently unclear how its induction contributes to disease. During natural infection, *Xam* enters through the stomata or other natural openings, multiplies in leaf tissues and then spreads systemically through the plant vascular system. It is not known whether induction of *MeSWEET10a* occurs in all tissue types or if it is limited to specific stages of infection. More generally, the spatial and temporal dynamics of *MeSWEET10a* induction during natural infections is unknown. To date, the tools required to address these questions have been lacking.

Gene editing technologies can facilitate the development of new tools for non-model organisms. Precise, error-free, knock-in products can be produced from the induction of DNA double-stranded break (DSB) repair pathways. This requires the introduction of a T-DNA that contains CRISPR/Cas9 elements plus a donor repair template. DSB repair is achieved by one of two pathways: either error-prone, non-homologous end-joining (NHEJ), or the more precise HDR pathway, with plant cells generally utilizing NHEJ (Puchta, 2005). HDR is well-established as an effective way to generate error-free products, but is extremely inefficient in plants, often occurring at rates less than 1% of transformed tissue (Fauser *et al*., 2012; Hahn *et al*., 2018; Paszkowski *et al*., 1988; Schmidt *et al*., 2019).

In this work, we developed a tool for visualizing CBB infection *in vivo*. This was done by exploiting the HDR pathway and the transcriptional activation of *MeSWEET10a* by TAL20 to generate cassava plants wherein *MeSWEET10a* was tagged with *GFP* in its native genetic context. More generally, a time-saving strategy was devised using a tissue-specific transcriptional activator designed to make the products of successful HDR identifiable early in the process of tissue culture. Using this strategy, we successfully selected and propagated edited plants to maturity. Furthermore, we re-activated expression of and visualized MeSWEET10a-GFP in regenerated plants, demonstrating their viability as tools for visualizing CBB infection. The strategy described herein could be widely applicable, and therefore of broad technical significance.

## RESULTS

### A time-saving strategy for identifying HDR products

A previous transcriptomics study revealed that *MeSWEET10a* expression is induced during CBB infection and required for full pathogen virulence (Cohn *et al*., 2014). However, the spatial and temporal dynamics of *MeSWEET10a* induction during this process remain unknown. We sought to develop a visual tool for monitoring CBB infection using CRISPR-based gene editing technology to tag *MeSWEET10a* with *GFP* at the endogenous locus. The endogenous expression pattern for *MeSWEET10a* does not lend itself to easy screening for HDR products, as the gene is normally only expressed in flowers (http://shiny.danforthcenter.org/cassava_atlas, Figures 1a, S1). Therefore, we developed the strategy outlined in Figure 1b. The T-DNA transformed into cassava callus tissue contained CRISPR components (the nuclease (Cas9) and gRNAs), a repair template (*GFP* flanked by homology arms), and a transcriptional activator (TA) component designed to facilitate screening early in the tissue culture process. The TA component consisted of a known transcriptional activator of *MeSWEET10a* from *Xam*, *TAL20*, driven by a tissue-specific promoter (from cassava gene *Manes.17G095200*) that is active primarily during the friable embryogenic callus (FEC) stage of cassava transformation and propagation (Figure 1c). Because ectopic expression and localization of MeSWEET10a-GFP at the plasma membrane may be detrimental, or even lethal during the tissue culture process, an Endoplasmic Reticulum (ER) -localization signal (HDEL) was included at the C-terminus of GFP in order to sequester the tagged protein (Gomord *et al*., 1997). The resulting construct (supplemental sequence file 170. pIICI_8469_17G095200pTAL20) carrying the above components was used to transform FEC derived from cassava cultivar TME 419.

**Figure 1.**
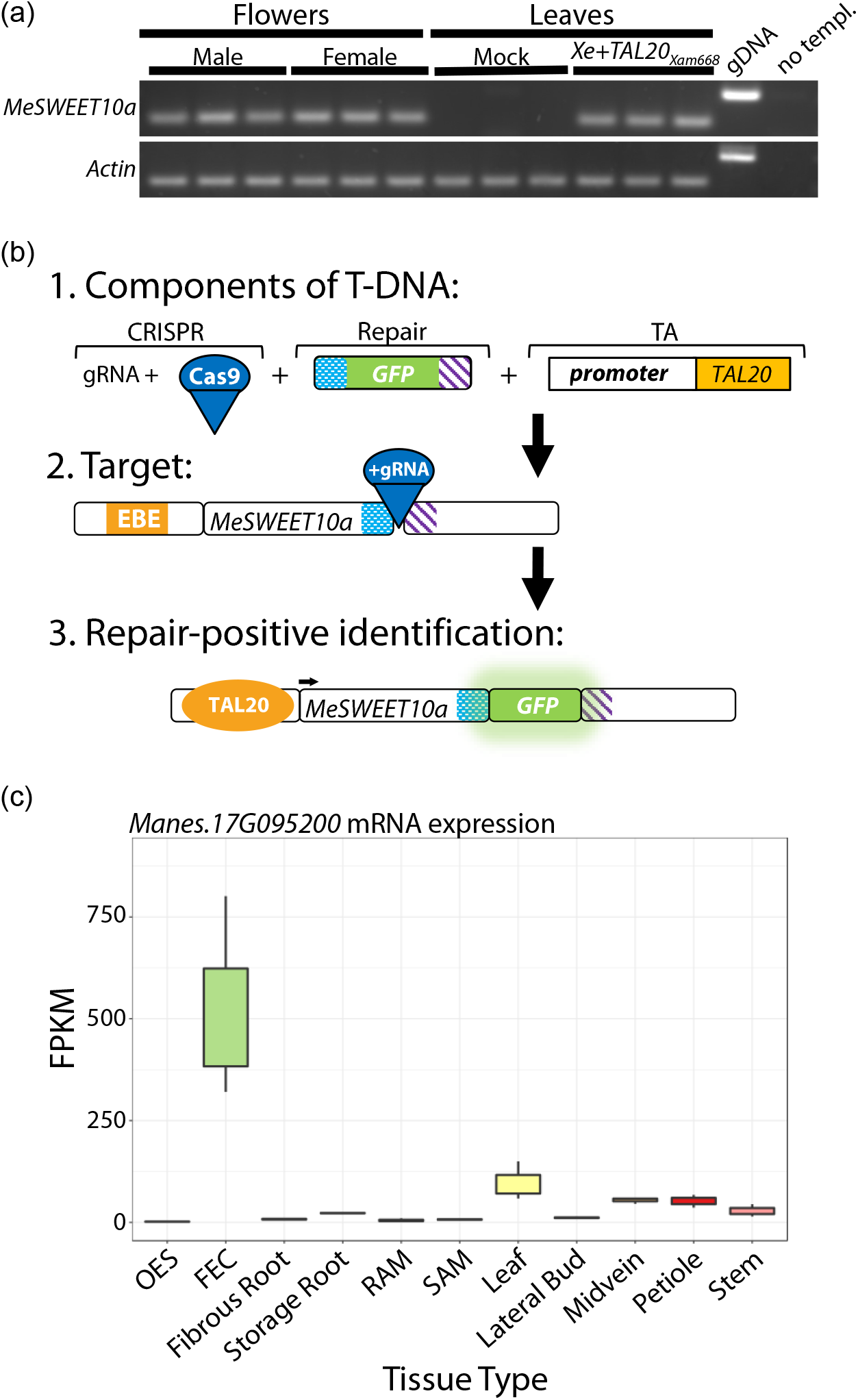
CRISPR-TA-coupled HDR strategy for tagging *MeSWEET10a*. **(a)** RT-PCR for *MeSWEET10a* expression in floral tissue. Analyses of male or female floral tissues are shown separately. *Xe*- and *Xe+TAL20_Xam668_*-infiltrated leaf tissues, harvested 2 days post-infiltration (dpi), were analyzed as negative or positive controls, respectively. Three biological replicates are presented for each tissue type. *Actin* was used as a loading control. Genomic DNA (gDNA) and no template controls are also shown. **(b)** Summary of the CRISPR-TA-coupled HDR strategy for tagging *MeSWEET10a*. Row 1: the supplied T-DNA includes CRISPR-based gene editing components (gRNA(s) and a nuclease, Cas9), a GFP-containing repair template (homology arms shown in blue/white and purple/white patterns), and a transcriptional activator (TA) system, consisting of a FEC tissue-specific promoter and the *TAL20* gene. Row 2: targeting the 3’ end of the *MeSWEET10a* locus. Cas9 + gRNAs cut just before the stop codon. Sequences homologous to the arms of the repair template are indicated in matched colored patterns. The Effector Binding Element (EBE) present in the promoter of *MeSWEET10a* is shown in orange. Row 3: Identification of repair-positive individuals using the TA system. TAL20 (orange circle) binds the EBE, activating *MeSWEET10a* expression (arrow). If the gene has been tagged in frame with *GFP*, fluorescence is detected through screening. Homology arms remain present (blue/white and purple/white patterns). **(c)** Expression pattern of gene from which the promoter was used in panel B. Y-axis: Fragments per Kilobase of transcript per Million mapped reads (FPKM) values. X-axis: tissue types included in tissue-specific RNA-seq data (Wilson *et al*., 2017). OES: organized embryogenic structures, FEC: friable embryogenic callus, SAM: shoot apical meristem, RAM: root apical meristem. Graph was produced using the available online Cassava Atlas tool (http://shiny.danforthcenter.org/cassava_atlas).

### Identification of HDR-positive lines

We used PCR to identify putatively successful products of HDR in which *GFP* is present at the 3’ end of *MeSWEET10a*. Genotyping plant tissue was carried out using a forward primer that binds the endogenous locus upstream of the LHA (left homology arm, i.e. not included in the repair template) and a reverse primer that either binds GFP or binds just downstream of the *MeSWEET10a* stop codon (Figure 2a). To estimate a base-line for HDR efficiency in cassava, we performed a large experiment wherein we sacrificed tissues early in the tissue culture process, 4 days after co-culture with Agrobacterium (Taylor *et al*., 2012). Healthy, actively proliferating embryogenic callus was used for DNA extraction. This tissue was subjected to antibiotic selection, but it was not screened for visual expression of GFP. In this case, each growing cluster of cells was considered an independent “line”, though it is not possible to know how many clusters would have subsequently developed to produce plants. Of the 300 lines selected, six were identified as positive for *GFP* insertion at *MeSWEET10a* using PCR (*MeSWEET10a* forward primer and *GFP* reverse primer), giving an estimate of HDR efficiency of 2% (Figure 2b). Because all transformed tissue was sacrificed for genotyping, no plant lines were recovered from this experiment.

**Figure 2.**
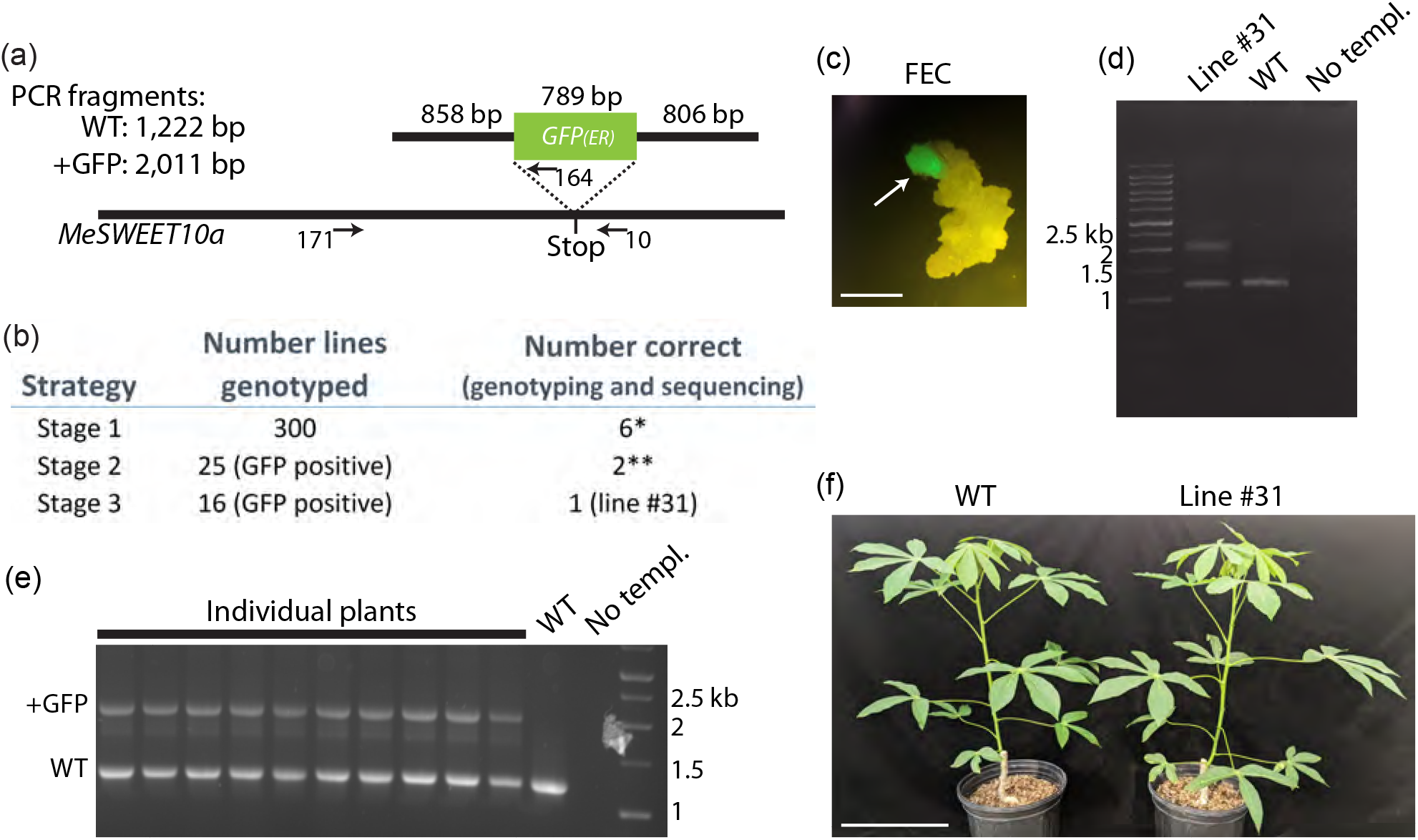
Genotyping and propagation of HDR-positive plants. **(a)** Schematic diagram illustrating the genotyping strategy at the *MeSWEET10a* locus (not drawn to scale). The repair-template with the sizes of the homology arms and *GFP* (with ER-localization signal) is shown above the gene locus. A dotted line illustrates the desired insertion site of *GFP* (just before the stop codon of *MeSWEET10a*). Arrows show approximate locations of primers used. Numbers correspond to primer sequences provided in Table S1. The sizes of PCR products using primers 171 and 10 are given with and without *GFP* integration. **(b)** Table summarizing genotyping from three separate experiments. Row 1: all growing FEC tissue was sacrificed just before stage 1 regeneration. No TA-based screening was performed. Rows 2 and 3: TA-based screening (GFP signal) performed for selection at either stage 2 or stage 3 regeneration. *All available tissue was used for genotyping. **Lines died in culture but were confirmed through genotyping and sequencing at the FEC stage (line 12 and 42, Figure S2). **(c)** Example image of epifluorescence signal used for screening GFP-positive FEC cells (arrow). Image taken just before stage 1. Scale bar = 1 mm. **(d)** Gel image of PCR products used to identify line #31. Primers 171 and 10 were used to genotype GFP-positive FEC tissue. Wild type (WT), no template (no templ.) controls are shown as well as a size standard with sizes in kilobases (kb) indicated. Bands from the positive line #31 were extracted from a gel and sequenced (Figure S2). **(e)** Same as panel D, but reaction was performed on leaf tissue from propagated plants. **(f)** Greenhouse-grown WT (left) and line #31 (right) adult plants. Scale bar = 25 cm.

Next, we investigated whether the TA system would increase likelihood of identifying successful HDR. Using the TA system coupled with antibiotic selection, FEC tissue was visually screened for a GFP signal five weeks after transformation, and healthy, proliferating GFP-positive tissues selected during either stage 2 or stage 3 of the process of somatic embryo regeneration (Figure 2c). Screening during stage 2 identified 25 GFP-positive lines. Of these, two were confirmed as successful products of HDR (lines 12 and 42, Figure 2b). Scarless insertion of *GFP* was validated by directly sequencing the PCR product generated using a *MeSWEET10a* forward primer and a *GFP*-specific reverse primer (Figure 2a, primers 171 and 164, Figure S2). Thus, the TA system successfully identified positive products of HDR. Unfortunately, the identified lines did not develop into viable plants. A subsequent round of transformation and GFP screening during stage 3 identified 30 GFP-expressing positive lines. Of these, line #31 was confirmed as a successful HDR event (Figure 2b, 2d). The two bands detected by PCR were consistent with a genotype hemizygous for the insertion of *GFP*. Subsequent PCR and sequencing confirmed successful, scarless insertion of *GFP* sequence at the 3’ end of *MeSWEET10a* in a hemizygous context (Figure S2). This result is consistent with the fact that successful HDR is relatively rare. However, in this case, the hemizygous nature of the insertion may have resulted from background heterozygosity at the gRNA target sites. We targeted the following sequences: gRNA1: CATTAACATTACCATTAACG and gRNA2: AGGGACACCAATGTCCTGCT. The gRNA2 spacer sequence is a perfect match for only one of the chromosomes, containing 2 mismatches (underlined) to the other copy of *MeSWEET10a*. As for gRNA1, resequencing the stock of TME 419 cassava plants used for this experiment at this locus found that the intended PAM site for gRNA1 (TGG) was not present (TGA) on either chromosome, thus this allelic site is not a target for gRNA1. Sequencing of the wild-type (WT) PCR band revealed no evidence of cutting in, or near, the targeted sequences, suggesting that the gRNAs may not be able to bind this allele (Figure S2). Plants were successfully regenerated from FEC of line #31, clonally propagated and established in the greenhouse. Line #31 plants were indistinguishable from wild-type (WT) plants, with no deleterious or “off-type” effects observed (Figures 2e, 2f). Thus, screening for the GFP signal enabled the identification of a viable, HDR-positive cassava plant line. Taken together, we conclude that the tissue-specific TA system, coupled with screening at the appropriate stage in tissue culture, facilitated in the identification of desired products of HDR. Specifically, this system was used to identify individuals harboring scarless integration of the *GFP* sequence at *MeSWEET10a*.

### MeSWEET10a-GFP as a tool for visualizing susceptibility gene expression

Ectopic expression of *MeSWEET10a* driven by TAL20 is necessary for full virulence of *Xam* (Cohn *et al*., 2014). Having tagged one of the endogenous copies of *MeSWEET10a* with *GFP*, we next tested whether expression of the fluorescently tagged protein would be induced when *Xanthomonas* delivered TAL20 into plant cells. Plants were infiltrated with either WT *Xam* or mutant *Xam* wherein TAL20 had been removed (*XamΔTAL20*) (Cohn *et al*., 2014). Additionally, TAL20 was introduced to another *Xanthomonas* species and was used to test for *MeSWEET10a* induction (*X. euvesicatoria* (*Xe+TAL20_Xam668_*)(Cohn *et al*., 2014). Consistent with previous reports, *MeSWEET10a* was detected in plants infiltrated with either WT *Xam* or *Xe+TAL20_Xam668_* (Figure 3a). To test for expression of the tagged product, a forward primer that binds outside of the LHA was paired with a *GFP* reverse primer and used for RT-PCR (to generate a 677 bp product). A product corresponding to *MeSWEET10a-GFP* was detected only in samples where TAL20 was present. No transcript was detected in WT plants. Confocal imaging was used to visualize fluorescent protein production in the presence of TAL20. GFP signal was detected in plants infiltrated with *Xe+TAL20_Xam668_* and not WT *Xe* (Figure 3b). The “spider-web” pattern observed in the GFP-positive sample is consistent with ER-localized signal (Nelson *et al*., 2007). This is expected, given the HDEL ER-localization signal present in the integrated *GFP* sequence (Gomord *et al*., 1997). The data demonstrates the viability of this strategy for the creation of a tool for visualization of the induction of *S* genes by pathogens and effectors *in vivo*. Generalizing the strategy could allow researchers to tag nearly any protein of interest in its native genomic context.

**Figure 3.**
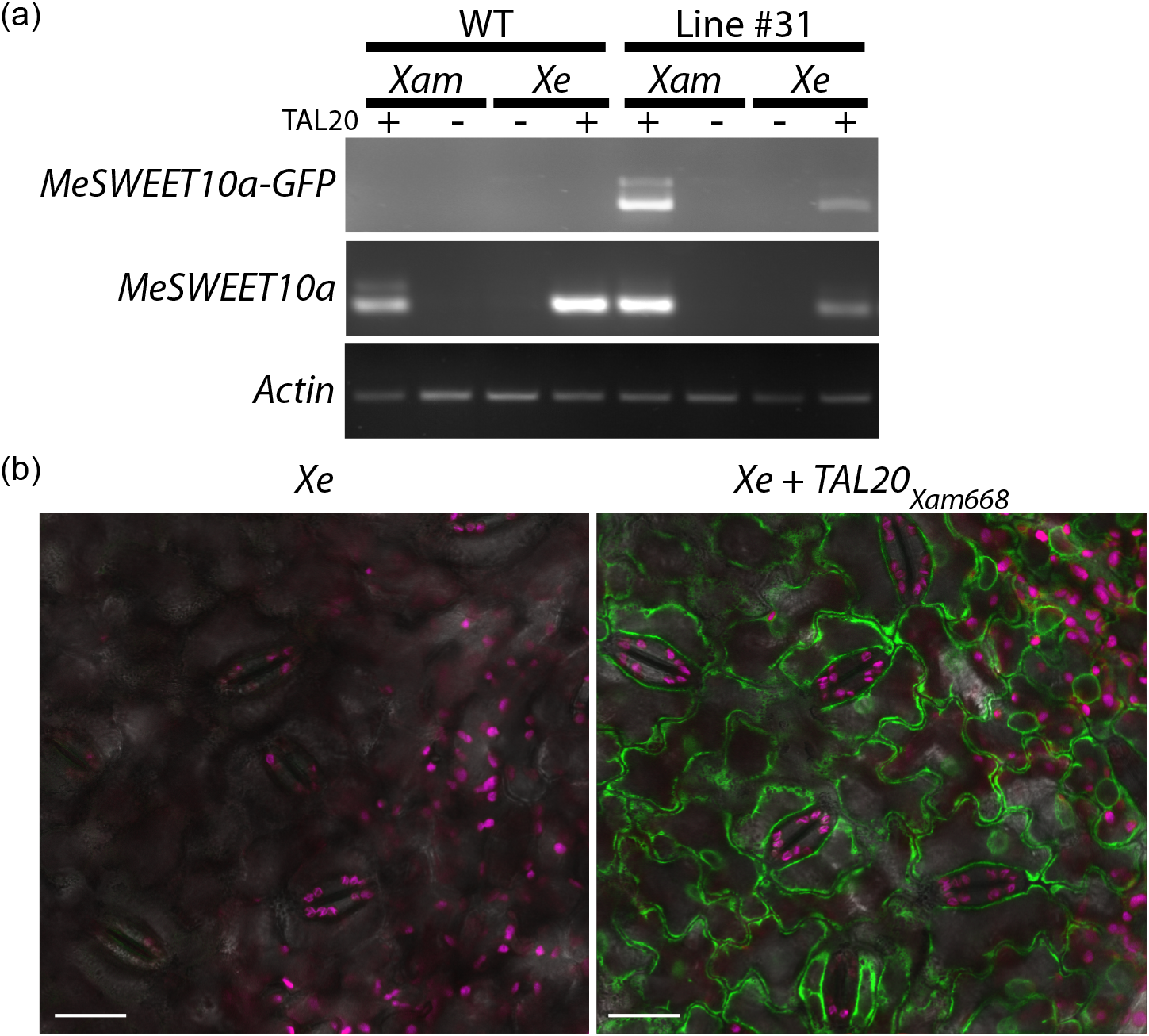
Activation of MeSWEET10a-GFP expression by *Xanthomonas*. **(a)** RT-PCR. Leaves from line #31 were infiltrated with the indicated *Xanthomonas* and RNA extracted at 3 dpi (3 biological replicates per sample). The presence (+) or absence (-) of TAL20 is indicated. Primers specific to *MeSWEET10a* with *GFP* were used for the first row and generic *MeSWEET10a* primers were used for the second row. *Actin* was used as a loading control. **(b)** Confocal laser scanning microscopy of the abaxial leaf epidermis of *Xanthomonas-* infiltrated leaves. Leaves from line #31 were infiltrated with either *Xe* or *Xe+TAL20_Xam668_* and imaged at 7 dpi. GFP is pseudocolored green, Chlorophyll – magenta, and bright field - grey. Scale bar = 20 μm.

## DISCUSSION

A major bottleneck exists in our knowledge regarding the process of infection by bacterial pathogens and how they manipulate host genes to promote disease at the molecular level. This knowledge will be essential to guide future efforts aimed at engineering robust, disease-resistant plants. For example, there are major gaps in our basic mechanistic understanding of the interface between the plant and the bacterial cell. Timing and location of effector delivery is not well understood. Are effector proteins capable of traveling cell-to-cell and/or do they act in ways that affect nearby cells? Are all cell layers within an infected plant organ (e.g. leaf, stem, root) subjected to the same infection strategy, and if so, are their responses varied? Is infection and subsequent effector delivery a constant process, or is it cyclical, happening in pulses, or in other patterns, possibly informed by environmental cues (e.g. circadian informed by light/dark cycles)? What happens when a plant encounters more than one pathogen at a time?

In this study, we used genome editing techniques to generate cassava plants that serve as tools for studying molecular-level CBB infection dynamics. A strategy was developed to facilitate the identification of successful products of HDR to produce plants harboring *MeSWEET10a* tagged with sequence encoding a fluorescent protein. This strategy resulted in scarless insertion of functional *GFP* at the endogenous locus of *MeSWEET10a*, allowing activation of MeSWEET10a-GFP expression by *Xam* and, more specifically TAL20_Xam668_, verified at the transcriptional and translational levels. These plants now serve as a valuable resource, enabling the visualization of *S* gene activation and CBB infection at the organ, tissue and molecular levels. Model plant systems are often used for tool development due to a relative abundance of resources. Though clearly valuable to the community, such findings or strategies often do not translate to crop systems and can be of limited use outside of the lab. Here, we describe a time-saving, viable approach, useful for answering the questions posed above about the mechanisms of plant-pathogen interactions. The work was performed in a non-model system that is also a farmer-preferred cultivar of cassava. Going forward, the development of such tools will be essential for answering detailed questions about disease processes and the timing of infection in this pathosystem. This strategy decreases the number of steps (and consequently the time) from technological development to its real-world impact.

In order to identify the rare, successful products of HDR, we developed an approach combining CRISPR-based genome editing technology with a tissue-specific TA system. It is worth noting that screening during the late stage of somatic embryo regeneration (stage 3, Figure 2b) resulted in the regeneration of viable, HDR-positive plants. Although we were able to identify HDR-positive lines earlier in the process of tissue culture, these lines did not reliably regenerate to become mature plants. The FEC TA system designed for this experiment also resulted in a number of false-positive lines when GFP screening was performed at stage 2 or stage 3. GFP-positive FEC tissue screened and selected at both stages 2 and 3 was not 100% the product of HDR (Figure 2b). Possible explanations for this include: 1) direct expression from the integrated T-DNA, if it had randomly inserted into an actively transcribed region of the genome, 2) false-negative genotyping by PCR (e.g. inefficient amplification or priming sites destroyed), or 3) spontaneous/unintended HR at a different locus that is actively transcribed. Despite efforts to identify these fluorescent products (by DNA-and RNA-based analysis), we observed no evidence to support the latter two possibilities, leaving expression from the randomly integrated T-DNA as the most likely explanation. Taken together, our data suggests that there is still much to learn about gene expression and the developmental reprogramming that occurs during the process of morphogenic tissue culture.

The TA-based HDR strategy outlined here presents a powerful approach for additional applications, especially for plant systems, in which HDR is a rare event (Schmidt *et al*., 2019). Given the time- and labor-intensive nature of cassava transformation, it is beneficial if successful products are identified early in the process of tissue culture. Because our gene of interest is not normally expressed in vegetative tissue, we chose a FEC-specific promoter driving a well-established transcriptional activator specific to this gene, TAL20_Xam668_. However, the true power inherent in this strategy is its adaptability to many other applications and systems, including the following: multiple and/or simultaneous knock-in events (where the rarity of HDR would be an even more limiting factor), epitope-tagging and allele swapping. Any HDR event for which screens or selection may be performed is compatible with this strategy, such as herbicide/antibiotic resistance or pigment production. Additionally, other tissue-specific or inducible TA systems can be employed via the use of an appropriate promoter (e.g. specific to seeds, cotyledons, hypocotyls, floral structures, etc.) coupled with either known or programmable transcriptional activators (e. g. dCas9-TV) (Li *et al*., 2017). Currently, certain applications of HDR in plant systems are not technically practical, and therefore not attempted. It is our hope that the work presented here demonstrates that given a well-designed approach, tools can be developed for the direct study of non-model systems.

## EXPERIMENTAL PROCEDURES

### Production of plants and growth conditions

Transgenic cassava lines of cultivar TME 419 were generated and maintained *in vitro* as described previously (Chauhan *et al*., 2015; Taylor *et al*., 2012). Briefly, transgenic FEC cells were selected for resistance using paramomycin (27.5 μM) on spread plates (also containing 125 mg/L cefotaxime). On stage 1, 2, and 3 plates, 45 μM paramomycin was used. Detection of GFP-derived fluorescence was assessed after three weeks in culture on stage 1, 2, or 3 regeneration media using a Nikon C15304 dissecting microscope equipped with an excitation filter of 460-500 nm and a barrier filter of 510 LP. Growth chamber settings used for tissue culture were 28°C +/- 1 °C; 75 μmol/m^2^/s light; 16 hrs light/8 hrs dark. In vitro plantlets were propagated and established in soil on a mist bench. Once established, plants were transferred to an open bench in the greenhouse with conditions of 28°C; humidity >50%; 16 hrs light/8 hrs dark; 1000 W light fixtures supplemented natural light levels below 400 W/m^2^.

### gRNA design and plasmids

Guide RNAs were designed to target the 3’ end of *MeSWEET10a* (gene identifier *Manes.06G123400, Manihot esculenta* genome v6.1). Sequences that consisted of a 5’-N_(20)_NGG-3’ motif were identified using the “Find CRISPR Sites” algorithm of Geneious, version 9.1.8 (Kearse *et al*., 2012). The software was used to exclude sites with potential off-target activity elsewhere in the cassava genome. The construct containing gRNAs (gRNA1: CATTAACATTACCATTAACG and gRNA2: AGGGACACCAATGTCCTGCT), Cas9 and the repair template was made using a modular system followed by final Golden Gate assembly as described previously (Čermák *et al*., 2017). Specifically, the final Golden Gate assembly consisted of three modules: pMOD_A0501 (encoding Cas9 and the Csy4 endoribonuclease), pMOD_B2103 (guide RNAs (gRNAs) separated by Csy4 spacers), pMOD_C0000 (the repair template), and a binary vector backbone (pTRANS_220d). The vector pMOD_A0501 was not modified. Guide RNAs were cloned into pMOD_B2103 using Golden Gate assembly following PCR synthesis of gRNA products using primers TC290-2, TC219, TC220, TC221, TC222 and TC222-2 (Table S1). The repair template (left homology arm (LHA, 866 bp), GFP (789 bp) and right homology arm (RHA, 815 bp)) was amplified using the following primer pairs: LHA (Bael-LHA-F1, Bael-LHA-R1) GFP (Bael-GFP-F2, Bael-GFP-R2) and RHA (Bael-RHA-F3, Bael-RHA-R3) and cloned into pMOD_C0000 using Gibson Assembly^®^ (New England BioLabs). Editing of the repair template by Cas9 was prevented by single base pair substitutions of the PAM sites, generating silent mutations in the coding sequence of the gene. All three final modules (A, B, C) were assembled into the backbone of the binary vector pTRANS_220d using Golden Gate cloning. The transcriptional activator system was then added to the resulting pTRANS_220d plasmid. The vector was linearized using SmaI and the activator system was added using In-Fusion cloning (In-Fusion^®^ HD Cloning Plus, Takara) according to manufacturer’s instructions. For the activator system, a tissue-specific promoter was identified (*Manes.17G095200*) showing high transcriptional activity in FEC cells (http://shiny.danforthcenter.org/cassava_atlas) generated by tissue-specific RNA-seq (Wilson *et al*., 2017). The Gene Finder tool in Cassava Atlas was used, with the search parameters “on” (FPKM > 300) in FEC, “off” (FPKM < 5) in organized embryogenic structures (OES), and “either” in all other tissue types. PCR (primers 17p-F and 17p-R) was used to amplify 615 bp of the promoter region upstream of the annotated transcriptional start site of the gene (*Manihot esculenta* genome v6.1, http://phytozome.jgi.doe.gov). *TAL20* from *Xanthomonas axonopodis* pv. *manihotis* (*Xam668*) was amplified by PCR (primers TAL20-F and TAL20-R) for use as the transcriptional activator of *MeSWEET10a*. The In-Fusion reaction was performed according to the manufacturer’s instructions. All primer sequences are listed in Table S1. The final construct was transformed into electrocompetent *Agrobacterium tumefaciens* strain LBA4404, and positive colonies selected on LB agar plates containing kanamycin (50 μg/ml), rifampicin (100 μg/ml) and streptomycin (30 μg/ml).

### Genotyping for HDR

Approximately 100 milligrams of tissue from either *in vitro*-grown plantlets or leaf tissue from greenhouse-grown plants was used for genomic DNA extraction (GenElute^™^ Plant Genomic DNA Miniprep Kit, Sigma). PCR was performed using primers SWEET10a_g-F and SWEET10aPro-R (Table S1). For Sanger sequencing, bands were isolated from a gel, subcloned (pCR4-TOPO TA Vector, Thermo Fisher Scientific) and sequenced (standard M13 primers) to verify zygosity and scarless-insertion.

### *MeSWEET10a* expression

#### *MeSWEET10-GFP* induction

*Xanthomonas* strains were grown on plates containing necessary antibiotics for 2 to 3 days at 30°C. The *Xanthomonas* strains used and their antibiotic resistances are as follows: 1. *Xam668* (rifampicin 50 μg/ml), 2.

*Xam668ΔTAL20* (suicide vector knockout, tetracycline 5 μg/ml, rifampicin 50 μg/ml) (Cohn *et al*., 2014), 3. *Xanthomonas euvesicatoria (Xe*, rifampicin 50 μg/ml) and 4. *Xe+TAL20_Xam668_* (kanamycin 50 μg/ml, rifampicin 50 μg/ml)(Cohn *et al*., 2014). Bacteria were scraped from the plates into 10 mM MgCl_2_ to concentrations of OD_600nm_ = 1. The first fully-expanded leaf of each plant was inoculated with a 1 ml needleless syringe using one bacterial strain per lobe and 5 to 10 injection points on each of 2 or 3 leaves. After inoculation, plants were kept under fluorescent light on a 12-hour day/night light cycle at 27°C for either 3 days (for RNA) or 7 days (for microscopy).

#### RNA

Three days post-infiltration (dpi), samples were taken using a 7 mm cork borer at each infiltration site, frozen in liquid nitrogen, ground to a fine powder and RNA was extracted (Spectrum^™^ Plant Total RNA Kit, Sigma). Biological replicates were pooled, such that each RNA sample contained multiple injection sites from 2 or 3 leaves. One microgram of DNase-treated RNA was converted to cDNA (SuperScript^™^ III Reverse Transcriptase, Invitrogen) and used for RT-PCR. Details of the PCR cycle numbers and primer sequences are shown in Table S1.

#### Confocal microscopy

leaves from plant line #31 were infiltrated with either *Xe* or *Xe+TAL20_Xam668_* and the abaxial side was imaged at 7 dpi. Square leaf samples of approximately 2 cm by 2 cm were taken near the infiltration point and mounted in a 35 mm Nunc coverglass bottom dish for microscopy. Confocal images were acquired at 1024×1024 pixel resolution on a Leica SP8-X using an HC PlanApochromat 63X Water Immersion Objective Lens with the pinhole set to 1 Airy unit. A 488 nm laser was used for excitation to image transmitted light with the spectral prism set to collect 493 to 585 nm and 654 to 779 nm emission wavelengths for GFP and chloroplast autofluorescence, respectively.

#### *MeSWEET10a* endogenous expression

Because WT cassava does not flower under our greenhouse conditions, flower tissue was collected from a CRISPR-generated line that has a mutation in the floral-repression pathway (*TERMINAL FLOWER1, TFL1*) (Adeyemo *et al*., 2019; Odipio *et al*., 2018). RNA was extracted (Spectrum^™^ Plant Total RNA Kit, Sigma) from male and female floral tissues. As a positive control for *MeSWEET10a* expression, WT cassava leaves were infiltrated with *Xe+TAL20_Xam668_* (Cohn *et al*., 2014) and RNA was extracted at 3 dpi and used for RT-PCR. Primers sequences are listed in Table S1.

## ACKNOWLEDGEMENTS

We are grateful for valuable advice and discussions with members of the Bart and Meyers labs as well as the DDPSC greenhouse staff for their work supporting this study. Additionally, we thank John Odipio for providing floral tissue for gene expression analysis. We acknowledge imaging support from Dr. Kirk Czymmek at the Advanced Bioimaging Center, Donald Danforth Plant Science Center, and usage of the Leica SPX-8 acquired through an NSF Major Research Instrumentation grant (DBI-1337680). This work was supported by the Bill and Melinda Gates Foundation (OPP1125410) and the National Science Foundation (NSF EDGE, award 1827761).

## AUTHOR CONTRIBUTIONS

K. V. Wrote manuscript, designed experiments and performed research / data analysis

I. O. Greatly contributed to experimental design and performed research

G. J. Performed research

M. Y. Performed research

N. T. Contributed to experimental design

B. C. M. Contributed to experimental design

R. B. Contributed to experimental design, data analysis, and manuscript writing

## SUPPLEMENTAL FILES

**Table S1.** Primers used in manuscript. (Supports all figures)

Each primer has a number and a name (first two columns). Column 3: the sequence of the primer (5’ to 3’ direction). Column 4: Description of each primer, how it was used, and any other relevant information.

**Figure S1.** Expression pattern of *MeSWEET10a* in vegetative tissues. (Supports Figure 1)

Y-axis: Fragments per Kilobase of transcript per Million mapped reads (FPKM) values. X-axis: tissue types included in tissue-specific RNA-seq data (Wilson *et al*., 2017). OES: organized embryogenic structures, FEC: friable embryogenic callus, SAM: shoot apical meristem, RAM: root apical meristem. Graph was produced using the available online Cassava Atlas tool (http://shiny.danforthcenter.org/cassava_atlas).

**Figure S2.** Sanger sequencing of successful knock-in lines. (Supports Figure 2)

Sanger sequencing of successful knock-in lines 31 (the focus of this manuscript), 12 and 42. Shown: raw sequencing data aligned to cultivar TME 419 (WT) reference sequence expected following scarless integration of GFP. The data are wrapped in multiple rows within a single page per sequencing product to aid in viewing. Page 1: sequencing data (base call and quality graphs) for line 31 from the upper (larger) band shown in the gel in Figure 2D (primers 171 and 10, Table S1). Page 2: data from the lower (smaller) band. Heterozygous SNPs are highlighted with colored backgrounds. These PCR products were sequenced using clone-seq with standard M13 sequencing primers. The products that were sequenced for lines 12 (page 3) and 42 (page 4) were generated using primers 171 and 164 (Table S1). Sequencing of these lines was performed directly on the PCR product using primers 200 and 172 (Table S1). Each page shows the sequencing data aligned to the expected product of HDR (highlighted in yellow). The LHA, GFP, and RHA sequence of the repair template are labeled in grey. Consensus sequence is given above the yellow-highlighted reference and sequencing data below. The names of the sequencing runs, which include the line number and either “large” or “small” for the bands sequenced from line 31 or the primer number used for lines 12 and 42, are to the left. Trimmed areas of poor sequence quality are indicated with a pink bar. Relevant primer sequences are shown in green. Analysis was performed using the algorithm built in to Geneious^®^ version 9.1.8.

