## Supplementary material for "Visualizing cassava bacterial blight at the molecular level using CRISPR-mediated homology-directed repair": Table S1

| Primer Number | Primer Name | Sequence (5'-3') | Description |
|---|---|---|---|
| 7 | TC290-2 | TGCTCTTCGCGCTGGCAGACATACTGTCCCAC | CmYLCV-F forward primer, type IIS site is highlighted in turquoise and overhang is underlined |
| 24 | TC219:GFPKIN-2 | TCGTCTCCGTAATGTTAATGCTGCCTATACGGCAGTGAACCTG | gRNA1-R reverse primer, type IIS site is highlighted in turquoise, gRNA spacer sequence green and overhang underlined |
| 25 | TC220:GFPKIN-2 | TCGTCTCATTACCATTAACGGTTTTAGAGCTAGAAATAGC | gRNA1-F reverse primer, type IIS site is highlighted in turquoise, gRNA spacer sequence green and overhang underlined |
| 26 | TC221:GFPKIN-2 | TCGTCTCCATTGGTGTCCCTCTGCCTATACGGCAGTGAAC | gRNA2-R reverse primer, type IIS site is highlighted in turquoise, gRNA spacer sequence green and overhang underlined |
| 27 | TC222:GFPKIN-2 | TCGTCTCACAATGTCCTGCTGTTTTAGAGCTAGAAATAGC | gRNA2-F reverse primer, type IIS site is highlighted in turquoise, gRNA spacer sequence green and overhang underlined |
| 12 | TC222-2 | TGCTCTTCTGACCTGCCTATACGGCAGTGAAC | 35Sterm-R reverse primer, type IIS site is highlighted in turquoise and overhang is underlined |
| 38 | Bael-LHA-F1 | CGCGTAGTCCTCGGTAGGCAAGCTTATTTAATTC | forward primer for the amplification of LHA flanked by 16bp of pMOD_C0000 for Gibson Assembly (blue) |
| 39 | Bael-LHA-R1 | GATTAGTCTTAACTTTGTCTGCAGG | reverse primer for the amplification of LHA flanked by 10bp of pMOD_C0000 for Gibson Assembly (blue) |
| 40 | Bael-GFP-F2 | AGACAAAGTTAAGACTAATCTTTTTCTCTTTCTCA | forward primer for the amplification of GFP flanked by 10bp of pMOD_C0000 for Gibson Assembly (blue) |
| 41 | Bael-GFP-R2 | GTTAATGTTATCAAAGCTCATCATGTTT | reverse primer for the amplification of GFP flanked by 10bp of pMOD_C0000 for Gibson Assembly (blue) |
| 42 | Bael-RHA-F3 | TGAGCTTTGATAACATTAACATTACCATTAACGTGA | forward primer for the amplification of RHA flanked by 10bp of pMOD_C0000 for Gibson Assembly (blue) |
| 43 | Bael-RHA-R3 | TGACTTGAAGTACACTCCGCTCTAGAATTAGTTG | reverse primer for the amplification of LHA flanked by 17bp of pMOD_C0000 for Gibson Assembly (blue) |
| 173 | 17p-F | TCGAGCTCGGTACCCACCCTACTTAAAAACCCTTTCG | forward primer for the amplification of Manes.17G095200 flanked by 15bp of pTRANS_220d for In-fusion cloning (blue) |
| 174 | 17p-R | ACGAATGGGATCCATGGTGACCTCTGGATTTCAGTACGAGG | reverse primer for the amplification of Manes.17G095200 flanked by 15bp of TAL20 product for In-fusion cloning (blue) |
| 175 | TAL20-F | ATGGATCCCATTCGTCCGCGCACGCCAAG | forward primer for the amplification of TAL20. First 15bp are included in Manes.17G095200 for In-fusion cloning (blue) |
| 176 | TAL20-R | GGCGCGCCGATCCCCTCACTGAGGAAATAGCT | reverse primer for the amplification of TAL20 flanked by 15bp of pTRANS_220d for In-fusion cloning (blue) |
| 171 | SWEET10a_g-F | CTTCCGTACGCGCTTCTTCATTTACTTTATATGCTAATAACTGACAAATACTG | forward primer used for genotyping products of HDR (Figure 2) |
| 10 | SWEET10aPro-R | CATAACCAAGAACACGTTAATGGTAATGTTAATG | reverse primer used for genotyping products of HDR (Figure 2) |
| 37 | Actin-F | ACAGTGTCTGGATCGGAGGATC | forward primer for Actin RT-PCR, 28 cycles |
| KV38 | Actin-R | GAAGCACTTCCTGTGGACGATG | reverse primer for Actin RT-PCR, 28 cycles |
| 200 | 10a_qPCR-F | CTTACTGTGTACCTTATCTATGCCACAAAGAAG | forward primer for MeSWEET10a and MeSWEET10a-GFP RT-PCR, 30 cycles |
| 201 | 10a_qPCR-R | GCCATGTGTAAGGAAAAGAGTTAAGATAGCG | reverse primer for MeSWEET10a RT-PCR, 30 cycles |
| 172 | GFP-R2 | CCAGTGAAAAGTTCTTCTCCTTTACTGAATTCGGCCG | reverse primer for MeSWEET10a-GFP RT-PCR, 30 cycles |
| 164 | GFP-R | CGTTGTGGGAGTTGTAGTTGTA | reverse primer for MeSWEET10a-GFP genotyping lines 12 and 42 (Figure S2) |
