## Supplementary figures and images for "Visualizing cassava bacterial blight at the molecular level using CRISPR-mediated homology-directed repair"

### Figure S1

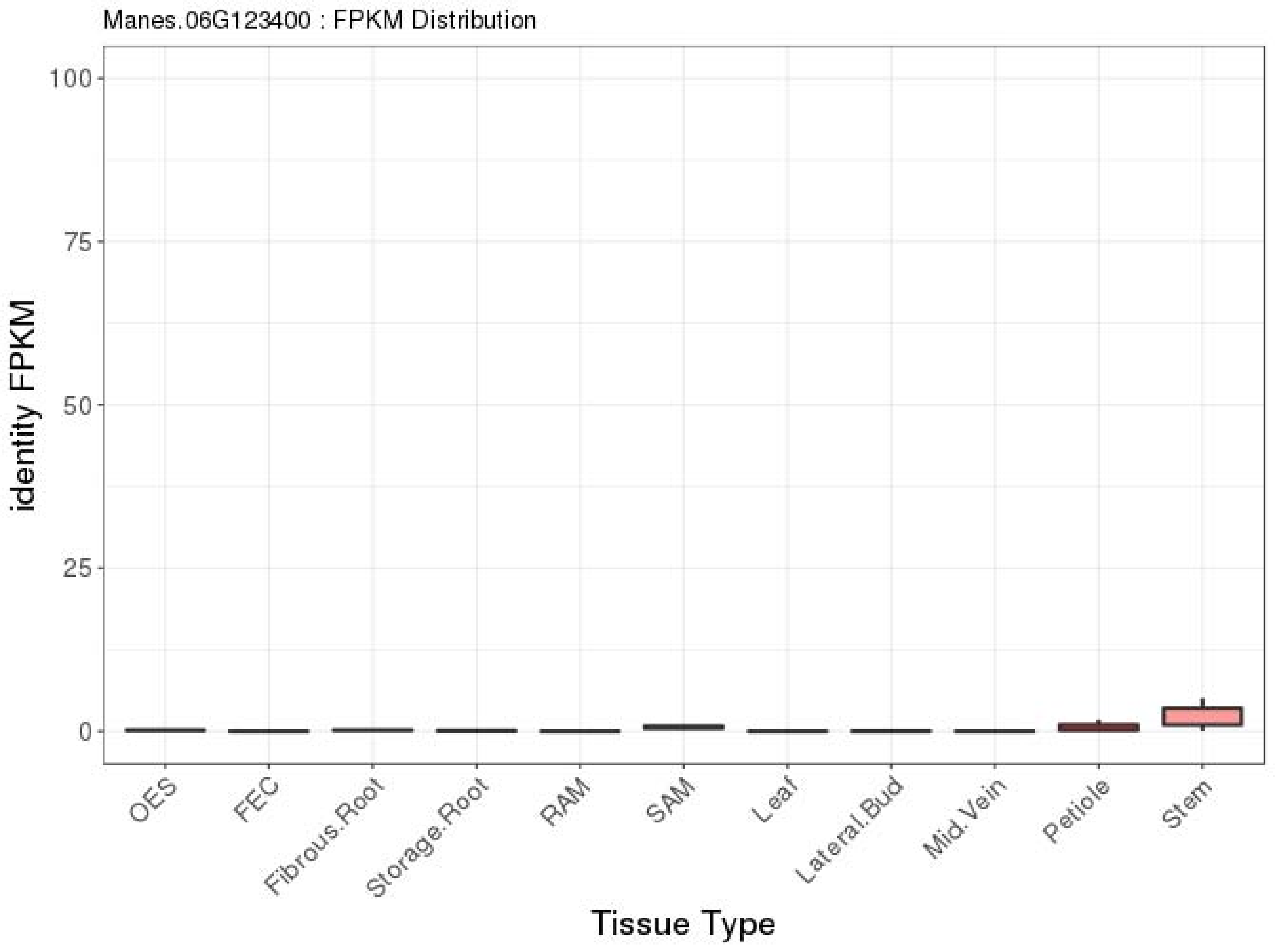

### Figure S2

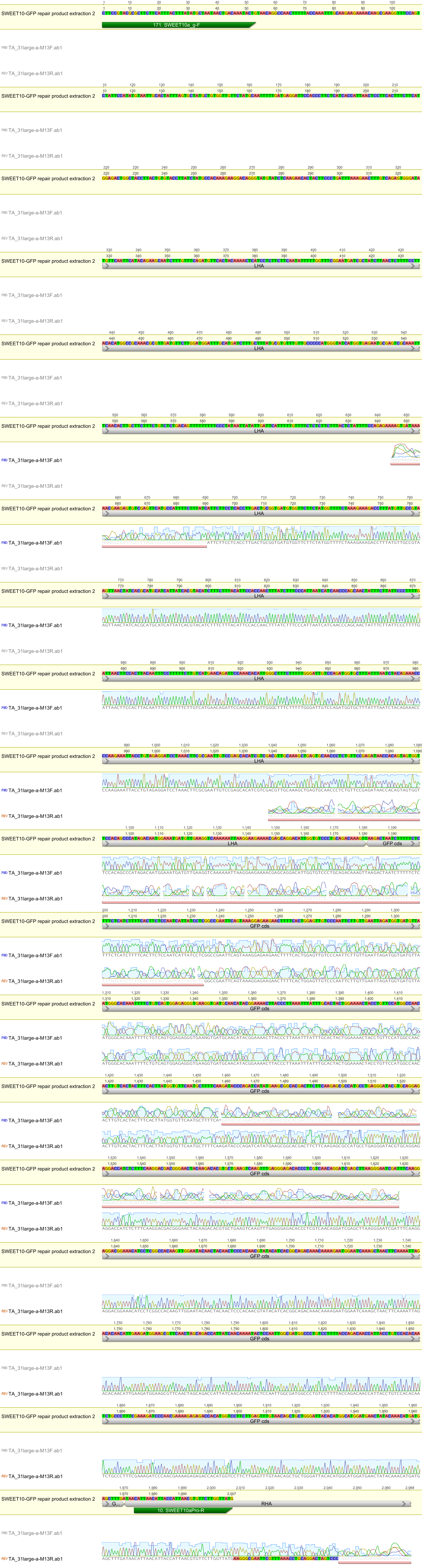

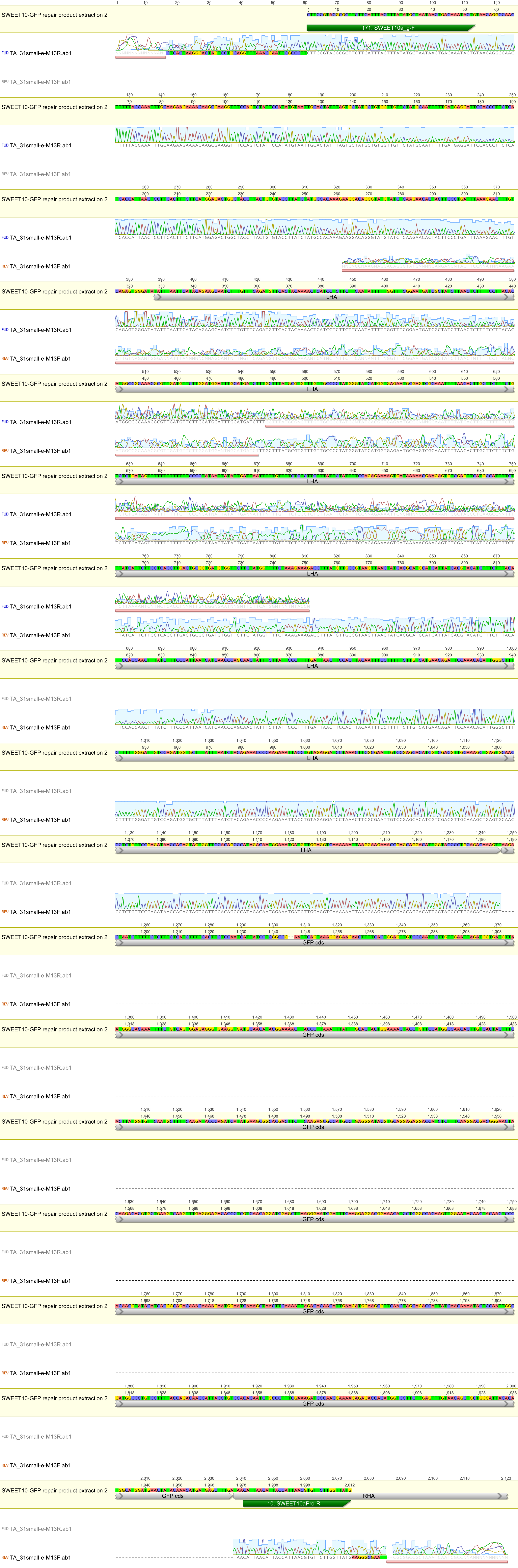

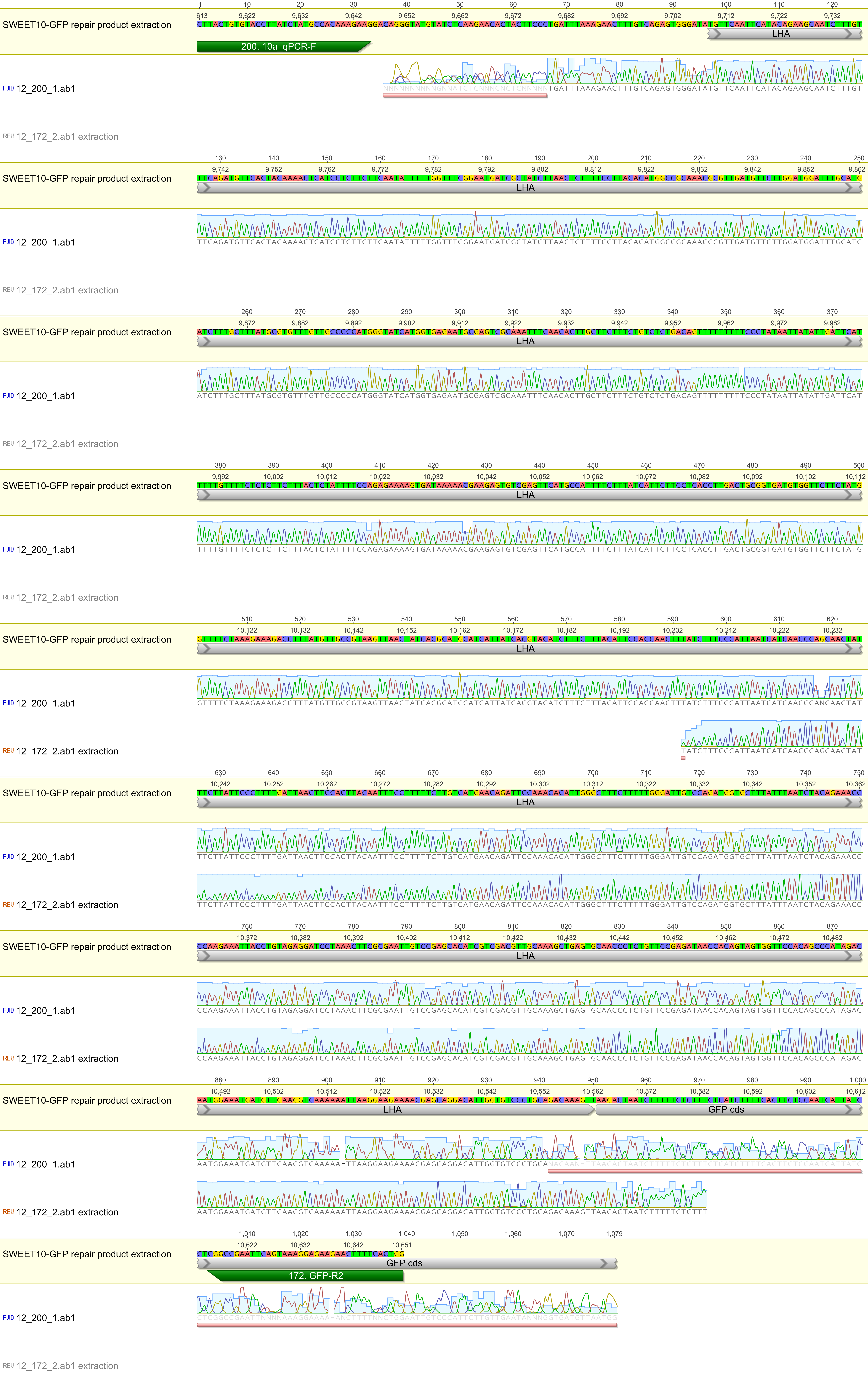

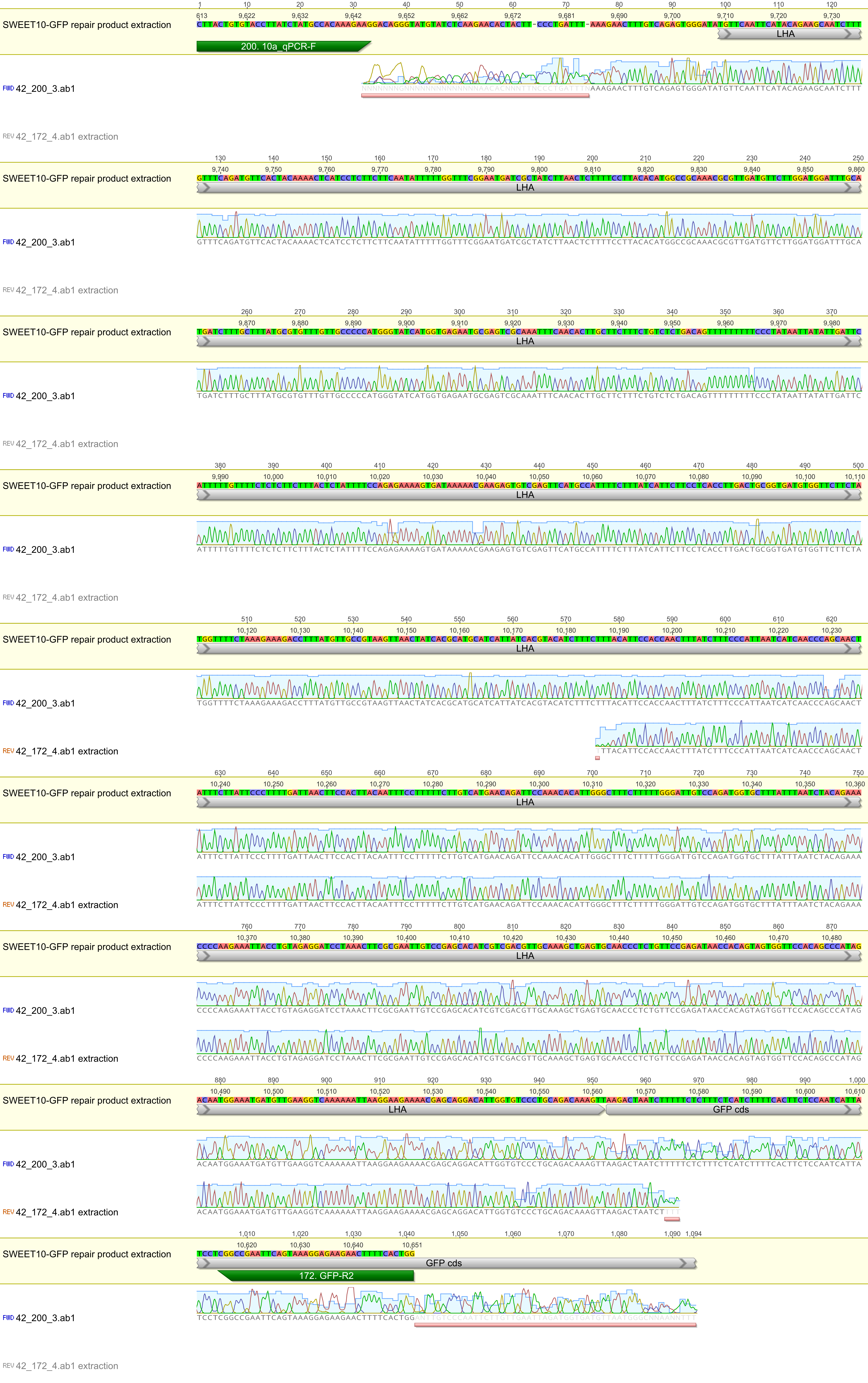
